# A comprehensive investigation of statistical and machine learning approaches for predicting complex human diseases on genomic variants

**DOI:** 10.1101/2022.05.16.492056

**Authors:** Chonghao Wang, Jing Zhang, Xin Zhou, Lu Zhang

## Abstract

**Background:** Quantifying an individual’s risk for common diseases is an important goal of precision health. The polygenic risk score (PRS), which aggregates multiple risk alleles of candidate diseases, has emerged as a standard approach for identifying high-risk individuals. A variety of tools have been developed to implement PRS. However, benchmarks for comparatively evaluating the performance of these different methods and for assessing their potential to guide future clinical applications are lacking.

**Results:** We systematically validated and compared thirteen statistical methods, five machine learning models and two ensemble models using simulated data, twenty-two common diseases with internal training sets and four diseases with external summary statistics from the UK Biobank resource. The effects of disease heritability, single nucleotide polymorphism (SNP) effect size and sample size are evaluated using simulated data. We also investigated the correlations between methods and their standard deviations of different diseases.

**Conclusions:** In general, statistical methods outperform machine learning models, and ensemble models, such as Super Learner, generally perform the best for most situations. We observed the correlations were relatively high if the methods were from the same category and the external summary statistics from large cohort GWAS could decrease the standard deviation of method correlations. By varying three factors in the simulated data, we also identified that disease heritability had a strong effect on the predictive performance of individual methods. Both the number and effect sizes of risk SNPs are important; and while sample size strongly influences the performance of machine learning models, but not statistical methods.

## Background

A key public health challenge is to identify individuals who are at high risk for common diseases, so as to facilitate prescreening or preventive therapies. Unlike rare diseases that are usually caused by inherited monogenic mutations, common diseases have multifactorial etiologies that involve the interplay of both genetic and non-genetic factors. Therefore, the effective identification of the risk factors contributing to the substantial burden of common diseases as well as high-risk individuals in the general population are core goals of precision health.

Rare variants with large effect sizes are recognized as good predictors of rare diseases such as Mendelian disorders. In common diseases, however, rare variants that confer several-fold increases in risks are usually absent from most of the general population. For example, 15 variants that occurred at significantly low frequencies in the general population were found to explain only 2% of coronary artery disease (CAD) heritability[1]. A recent study also showed that the strongest signal in the *LDLR* gene, which conferred a fourfold increased risk for myocardial infarction, could only be identified in 2% of patients[2]. Thus, strategies that leverage only rare variants are likely to achieve limited success in predicting the common disease risks. In contrast, with many common risk variants identified in genome-wide association studies (GWAS), the polygenic risk score (PRS) has emerged as a promising approach for using common variants to predict disease risks[3-5]. The PRS calculates an individual’s genetic risk for a given disease by aggregating the common risk alleles (minor allele frequency > 1%) that the individual carries, weighted by their effect sizes, that is, the logarithms of their odds ratios (ORs). The performance of a PRS could be influenced substantially by the sample size of the GWAS, and also by the number of single nucleotide polymorphisms (SNPs) involved. For example, a study using a multilocus PRS involving 13 genome-wide significant SNPs (p < 5 × 10^−8^) identified only 20% of individuals in the healthy population who had 1.4 times or greater risks for CAD[6], demonstrating the limited predictive power of approaches using only genome-wide significant SNPs. The low predictive power of the approaches that only aggregate genome-wide significant risk alleles motivated us to optimize PRS calculation by selecting the most effective SNPs with accurate effect sizes. Many methods for calculating the PRS have been developed to deal with this limitation; these methods can be classified under four broad categories: three types of statistical methods, including thresholding-based methods, Bayesian-based methods, and penalized regression methods; as well as machine learning models (Table 1).

**Table 1.**
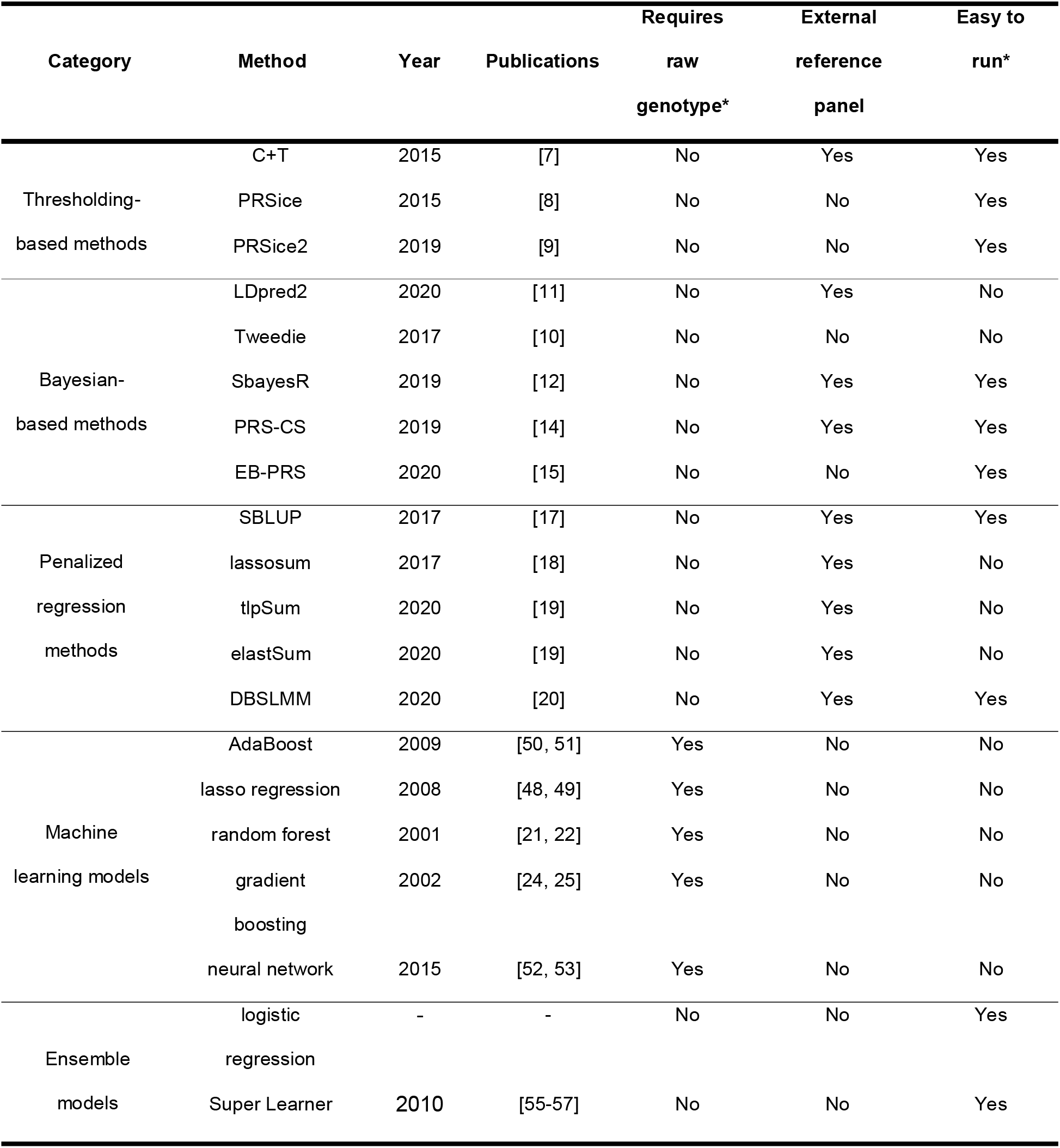
Overview of the statistical methods and machine learning models evaluated in the study. The command lines for each method are described in the Additional file 13: Supplementary Note.

Thresholding-based methods are designed to explore the sophisticated thresholds of SNP p-values, and to remove all redundant association signals; they achieve this by considering the pairwise linkage disequilibrium (LD) between SNPs. Conventional thresholding-based methods randomly “prune” SNPs according to their pairwise LD, before selecting the eligible SNPs with p-values lower than the given threshold; this strategy is commonly referred to as pruning plus thresholding (P+T). “LD clumping” (C+T)[7] improves P+T by removing less significant SNPs iteratively and keeping the more significant SNPs in every iteration. The PRS of C+T is then calculated by aggregating the remaining *n* SNPs according to their regression coefficients *β*_*i*_ (from logistic regressions that include only the target SNPs and covariates as predictors) and the copies (*S*_*i*_) of effect alleles, as follows: 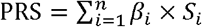. PRSice[8] extends C+T method and automatically examines a large number of p-value thresholds, then selects the most powerful threshold after correcting for population stratification and clumping the linked SNPs. PRSice2[9], which optimizes the workflow of PRSice by reducing demands on memory and speeding up PRS calculations, is better suited to Biobank-scale data.

Instead of removing linked SNPs with marginal p-values, Bayesian-based methods correct SNP effect sizes by considering the neighboring LD patterns; they also consider all SNPs during PRS calculation. LDpred[7] designs a Bayesian model to estimate posterior mean effect sizes from the summary statistics of the GWAS as well as LD patterns from an external reference panel. Tweedie[10] implements an ultra-fast empirical Bayes estimation for SNP effect sizes without assuming any prior distributions, thus avoiding the need for parameter tuning. LDpred2[11] resolves dedicated issues encountered in LDpred; it estimates the proportion of causal variants more efficiently and fine-tunes hyperparameters using two new functions: “sparse” and “auto”. SbayesR[12] extends Bayesian multiple regression[13] to accommodate GWAS summary statistics and adopts a finite mixture of normal distributions prior to perform a Bayesian posterior inference on the effect sizes of SNPs. PRS-CS[14] incorporates continuous shrinkage priors to estimate SNP effect sizes in a Bayesian regression framework; by doing so, it improves over LDpred and SbayesR, which suffer from problems with inaccurate local LD adjustment due to the discrete mixture priors. Unlike other Bayesian-based methods, EB-PRS[15] infers the SNP LD distributions from the input data directly, instead of relying on external reference panels.

Penalized regression methods jointly consider the effect sizes of different SNPs while penalizing the SNPs with small effect sizes in the prediction. BLUP[16] and SBLUP[17] are conventional algorithms to estimate individuals’ total genetic effects as random effects in mixed linear models. Lassosum[18] calculates PRS by solving a convex optimization problem of penalized regression and avoids the nonconvergence issue encountered in LDpred. Lassosum also includes a strategy to fine-tune parameters in the absence of a validation set. penRegSum[19] improves on lassosum by using a “truncated lasso penalty” and an “elastic net” to encourage sparsity and handle covariates more efficiently. DBSLMM[20] implements a deterministic Bayesian sparse linear mixed model, which fits a linear mixed model using GWAS summary statistics, LD matrix and the LD block information. DBSLMM relies on a simple deterministic search algorithm to achieve more efficient analyses with Biobank-scale data.

Besides the abovementioned three types of statistical methods, several machine learning models have been applied to predict disease susceptibility by supervised learning. Random forest[21, 22] is an ensemble method that trains a multitude of decision trees using sub-sampled training data. Chung et al.[23] identified 289 candidate markers with good discriminability and achieved an area under the receiver operating characteristic curve (AUROC) of 0.702 to predict individuals’ susceptibility to bipolar disorder using random forest. Gradient boosting[24, 25] is also an ensemble model, which trains decision trees based on residuals instead of assigning weights to different stumps. A study on breast cancer in two Finnish cohorts applied gradient boosting[26] to model complex SNP–SNP interactions and found this to be a better predictor than the PRS. GraBLD[27] includes gradient-boosted regression trees to correct SNP effect sizes and implements a regional adjustment for LD correction; its performance was shown to be comparable to LDpred when applied to Biobank-scale GWAS data. Romagnoni et al.[28] compared the performance of common machine learning models in classifying Crohn’s disease patients and found that nonlinear models such as gradient-boosted trees and neural networks were more robust than penalized logistic regressions. Deep neural network[29], which considers the non-additive effects of SNPs, has also been proposed as a promising technology to improve disease prediction. For instance, by implementing a fully connected neural network, DL-PRS[30] learns the weight of each SNP, before transferring them to the hidden layers of the neural network. It achieved outstanding improvements in predicting chronic obstructive pulmonary disease with data from the UK Biobank.

The statistical methods and machine learning models presented above have demonstrated substantial power for disease prediction in a variety of studies. However, a benchmark for evaluating their respective performance in providing practical guidance for clinical applications is lacking. In this study, we performed a comparative analysis of the performance of thirteen statistical methods, five machine learning models and two ensemble models for disease prediction (Table 1) on simulated data, twenty-two common diseases (*D*_22_) with internal training sets and four common diseases (*D*_4_) with external summary statistics from UK Biobank (**Methods**). Their performance was evaluated by AUROC, the value of the area under the precision-recall curve (AUPRC), Pearson correlation coefficient (PCC) and the risk in the ascending percentile of prediction scores (odds ratios, ORs, **Methods**). We identified the sophisticated ensemble models were the best choices for most situations. We also discussed the effects of disease heritability, SNP effect sizes and sample sizes on disease predictions, calculated the correlations between different models and compared the relative suitability of statistical methods and machine learning models to different scenarios. The analytical procedure used in this study is illustrated in Figure 1.

**Figure 1.** Workflow for evaluating the statistical methods, machine learning models and ensemble models for disease prediction on simulated and UK Biobank data.

## Results

### Methods comparison using simulated data: influence of disease heritability

We observed that the predictive power of all methods benefited from high disease heritability 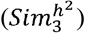, owing to the fact that greater phenotype variations can be explained by the genetic variations (Table 2, Figure 2A and 2B). When disease heritability is low 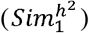, the performance of all statistical methods is poor; however, they still outperform the machine learning models. When disease heritability is approximately 0.8 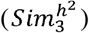, thresholding-based methods perform the best (average AUROC = 0.91, average AUPRC = 0.91 and average PCC = 0.68); only EB-PRS, a Bayesian-based method, achieves comparable performance (AUROC = 0.91, AUPRC = 0.91 and PCC=0.69). Interestingly, we noticed that the “simplest” thresholding-based method—C+T—performed the best overall. This is probably because the small proportion of SNPs with the strongest signals provides sufficient power to distinguish cases and controls, while LD thresholding removes noise from redundant signals. In general, different statistical methods achieve relatively stable AUROC, AUPRC and PCC; the exception is the penalized regression method, SBLUP, at 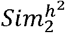 and 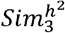 (Figure 2A, 2B and Additional file 1: Figure S1A). In contrast, the performance of machine learning models is poor overall, even when disease heritability increases to 0.8 (average AUROC = 0.73, average AUPRC = 0.71 and average PCC=0.40).

**Table 2.** Parameters for the simulated data with respect to disease heritability (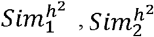 and 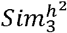) SNP effect sizes 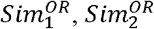 and 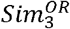) and sample sizes (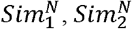 and 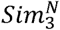)

| Simulated<br>datasets | Cases/Control<br>s in training<br>set (in<br>thousands) | Cases/Control<br>s in validation<br>set (in<br>thousands) | Cases/Contr<br>ols in test set<br>(in<br>thousands) | No. of causal SNPs with<br>effect sizes of<br>Small/Moderate/Large | $h^2$ |
| --- | --- | --- | --- | --- | --- |
| $Sim_1^{h^2}$ | 20/20 | 10/10 | 10/10 | 2,500/5,000/2,500 | 0.2 |
| $Sim_2^{h^2}$ | 20/20 | 10/10 | 10/10 | 2,500/5,000/2,500 | 0.5 |
| $Sim_3^{h^2}$ | 20/20 | 10/10 | 10/10 | 2,500/5,000/2,500 | 0.8 |
| $Sim_1^{OR}$ | 20/20 | 10/10 | 10/10 | 1,000/0/0 | 0.5 |
| $Sim_2^{OR}$ | 20/20 | 10/10 | 10/10 | 0/1,000/0 | 0.5 |
| $Sim_3^{OR}$ | 20/20 | 10/10 | 10/10 | 0/0/1,000 | 0.5 |
| $Sim_1^N$ | 5/5 | 2.5/2.5 | 2.5/2.5 | 2,500/5,000/2,500 | 0.5 |
| $Sim_2^N$ | 10/10 | 5/5 | 5/5 | 2,500/5,000/2,500 | 0.5 |
| $Sim_3^N$ | 30/30 | 15/15 | 15/15 | 2,500/5,000/2,500 | 0.5 |

**Figure 2.** AUROC and AUPRC values of the method results on simulated data by varying disease heritability (**A** and **B**), SNP effect sizes (**C** and **D**) and sample sizes (**E** and **F**).

Another crucial component of a method for disease prediction (apart from general predictive power) is the capacity to identify high-risk individuals in the general population. We calculated ORs for the individuals with the top 1, 5, 10 and 20 percentiles of PRSs by comparing the odds of individuals with the prediction scores that were higher or lower than the corresponding thresholds (**Methods**). All methods perform poorly in identifying high-risk individuals at 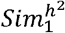, even for the individuals possessing the top 1 percentile of PRS (average OR = 5.4). The ORs become greater with increasing disease heritability and with decreasing the target thresholds (Figure 3A). The trends in the ORs of the different methods generally reflect the ones shown in their AUROCs, AUPRCs and PCCs. Thresholding-based methods outperform other methods for individuals with PRS above the top 5 percentile, with C+T even achieving outstanding power in the top 10 percentile. The machine learning models perform relatively poorly in comparison with most statistical methods. Our results suggest that the PRS is only effective for predicting diseases with high heritability and where individuals within the top 5 percentile are deemed to be at high risk.

**Figure 3.**
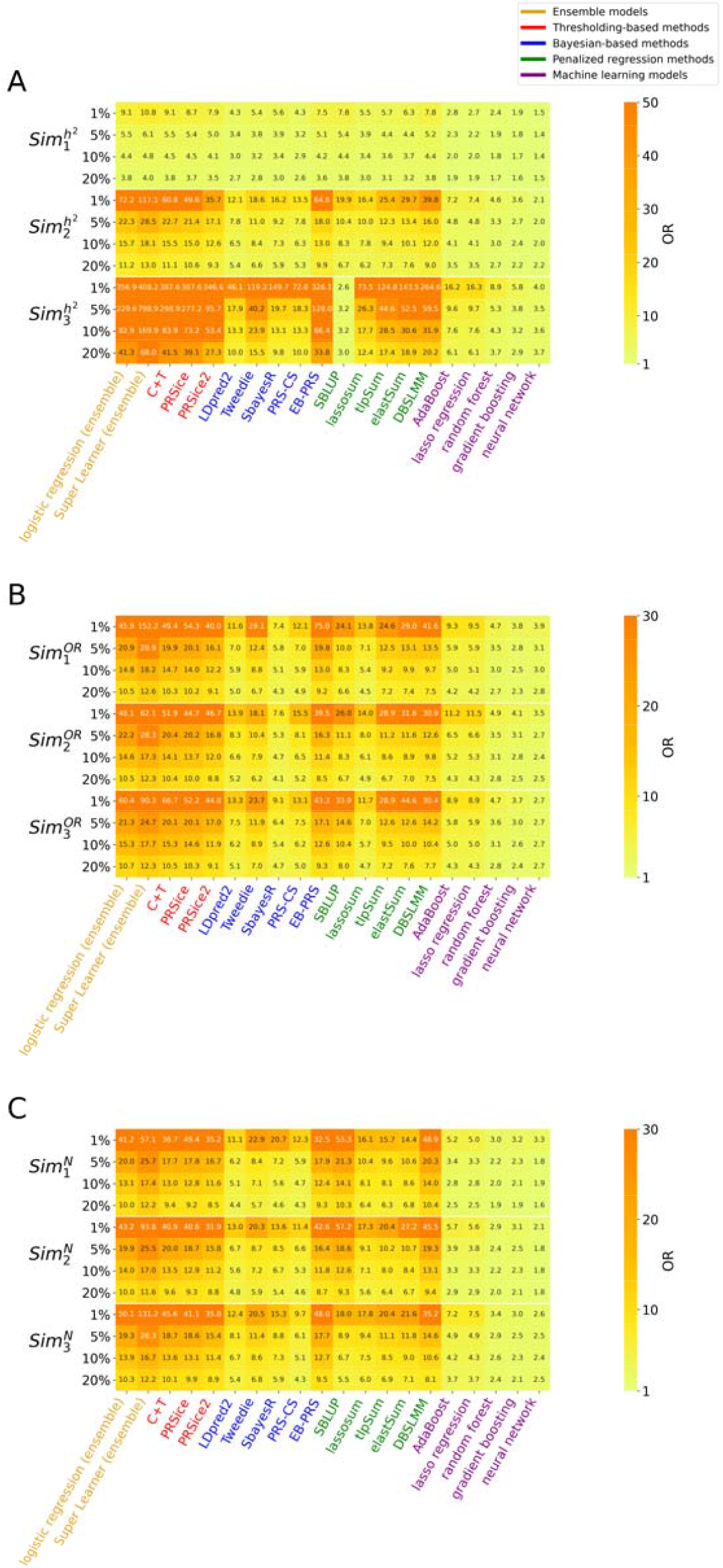
The ORs for the top 1, 5, 10 and 20 percentiles of the method results on the simulated data by varying disease heritability (**A**), SNP effect sizes (**B**) and sample sizes (**C**).

### Methods comparison using simulated data: influence of SNP effect sizes

Across all methods, predictive performance is not strongly affected by the variation in SNP effect sizes (Table 2, Figure 2C, 2D and Additional file 1: Figure S1B). It is conceivable that although individual SNPs with small effect sizes 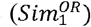) would have made it almost impossible to distinguish cases and controls, the sufficiently large number of such SNPs (1,000 in our experiments) could have allowed for significant improvements in performance that is comparable to datasets with large SNP effect sizes 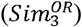.

Thresholding-based methods perform better than others (average AUROC=0.82, average AUPRC=0.81 and average PCC = 0.55). Among the Bayesian-based methods and penalized regression methods, EB-PRS (average AUROC=0.81, average AUPRC=0.80 and average PCC = 0.54) is comparable to PRSice2, while the performance of SBLUP, DBSLMM, elastSum and tlpSum is slightly poorer than that of thresholding-based methods. All machine learning models perform more poorly than statistical methods. There are no significant changes in the ORs of the top percentiles across different SNP effect sizes. The methods achieving the top AUROCs and AUPRCs also have outstanding abilities for identifying high-risk individuals in general populations. The absolute OR values of machine learning models, although acceptable (their ORs ranged from 2.3 to 11.5), are still poorer than those achieved by statistical methods (Figure 3B).

### Methods comparison using simulated data: influence of sample size

In general, the three thresholding-based methods (C+T, PRSice and PRSice2), as well as the EB-PRS and DBSLMM, achieve good predictive performance regardless of the numbers of samples that are used for model fitting (Table 2, Figure 2E, 2F and Additional file 1: Figure S1C). SBLUP only achieves good predictive performance at a small sample size 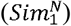). We found that an increase in sample size had different effects on the method performance. Among thresholding-based methods, a mild shift in predictive power is observed with increasing sample size. With the exception of PRS-CS, the same trend is observed among Bayesian-based methods; it is also apparent here that a larger sample size is beneficial to disease prediction. In contrast, among penalized regression methods, SBLUP and DBSLMM perform better when sample sizes are small; both two methods perform best at 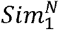. Although different trends with varying sample size are observed among the statistical methods, the standard deviations of their AUROCs, AUPRCs and PCCs generally decrease with increasing sample size. While the machine learning models perform poorly using the simulated datasets, their performance improves with increasing sample size. The OR patterns of statistical methods are broadly consistent with those of the AUROC, AUPRC and PCC. However, among the machine learning models, neural network and gradient boosting show opposite trends in ORs compared with those of the AUROC, AUPRC and PCC (Figure 3C). Although the general performance of neural network and gradient boosting improves with increasing sample size, they fail to achieve substantial increases in ORs.

### Methods comparison using real data: predicting twenty-two common diseases with internal training data

We used the SNP genotypes of 488,377 participants from the UK Biobank cohort, who were aged between 42 and 69 years old during the baseline examinations (in years between 2006 and 2010). After applying quality control, we extracted prevalent and incident cases from 337,536 unrelated white British participants on the basis of their self-reports and hospitalization records (**Methods**). Twenty-two common diseases with at least 6,000 prevalent cases and 100 incident cases were chosen for our investigation (*D*_22_, Additional file 6: Table S1). We split the prevalent cases into two parts: two-thirds of the cases were used in the training set and the remaining one-third was used for the validation set. The incident cases were used as a test set to evaluate a model’s predictive performance. A pooled dataset comprising 93,499 controls (45,872 men and 47,627 women) was assembled for participants who were never diagnosed for any of the twenty-two diseases; these participants were randomly selected to be controls for training, validation and test sets (**Methods**).

We found that the AUROCs, AUPRCs and PCCs of all methods were slightly better than random for most of the selected diseases (Additional file 2: Figure S2A, S2B and Additional file 3: Figure S3A). Two diseases, diabetes and asthma, achieve the highest average AUROC and AUPRC, at values above 0.6 (Figure 4A and 4B). With the exception of SbayesR and neural network, all methods perform comparably well for predicting diabetes, and thresholding-based methods perform the best (Figure 4A). Except neural network (AUROC = 0.64, AUPRC = 0.63 and PCC = 0.25), the performance of machine learning models is poor for asthma prediction (Figure 4A, 4B and Additional file 3: Figure S3A), while that of the statistical methods are comparable, such as SbayesR (AUROC = 0.63, AUPRC=0.61 and PCC = 0.22), PRS-CS (AUROC = 0.62, AUPRC = 0.59 and PCC = 0.19), tlpSum (AUROC = 0.62, AUPRC = 0.61, PCC=0.21), elastSum (AUROC = 0.62, AUPRC=0.60 and PCC=0.21). The ORs of top percentiles are not convincing and their trends of some diseases are even contradictory to expectations (Figure 5A and Additional file 4: Figure S4A). For example, the ORs of DBSLMM are 0.2 (1%), 1.0 (5%), 1.4 (10%) and 2.1 (20%) for asthma. We investigated this phenomenon and found that these questionable values arose from instances where there were few incident cases. The incident cases in the UK Biobank were derived from hospital records, which applied more stringent criteria for identifying disease than self-reported cases (Additional file 6: Table S1). To verify the performance of the disease prediction methods on a larger dataset, we performed a second round of analysis using a workflow that was modified slightly from that shown in Figure 1. In the new analysis, we used 2/9 of the prevalent cases for the validation sets—instead of 1/3 as before— and used 1/9 of the prevalent cases as new test sets. These test individuals are not accessible to the model during training or hyperparameter selection.

**Figure 4.**
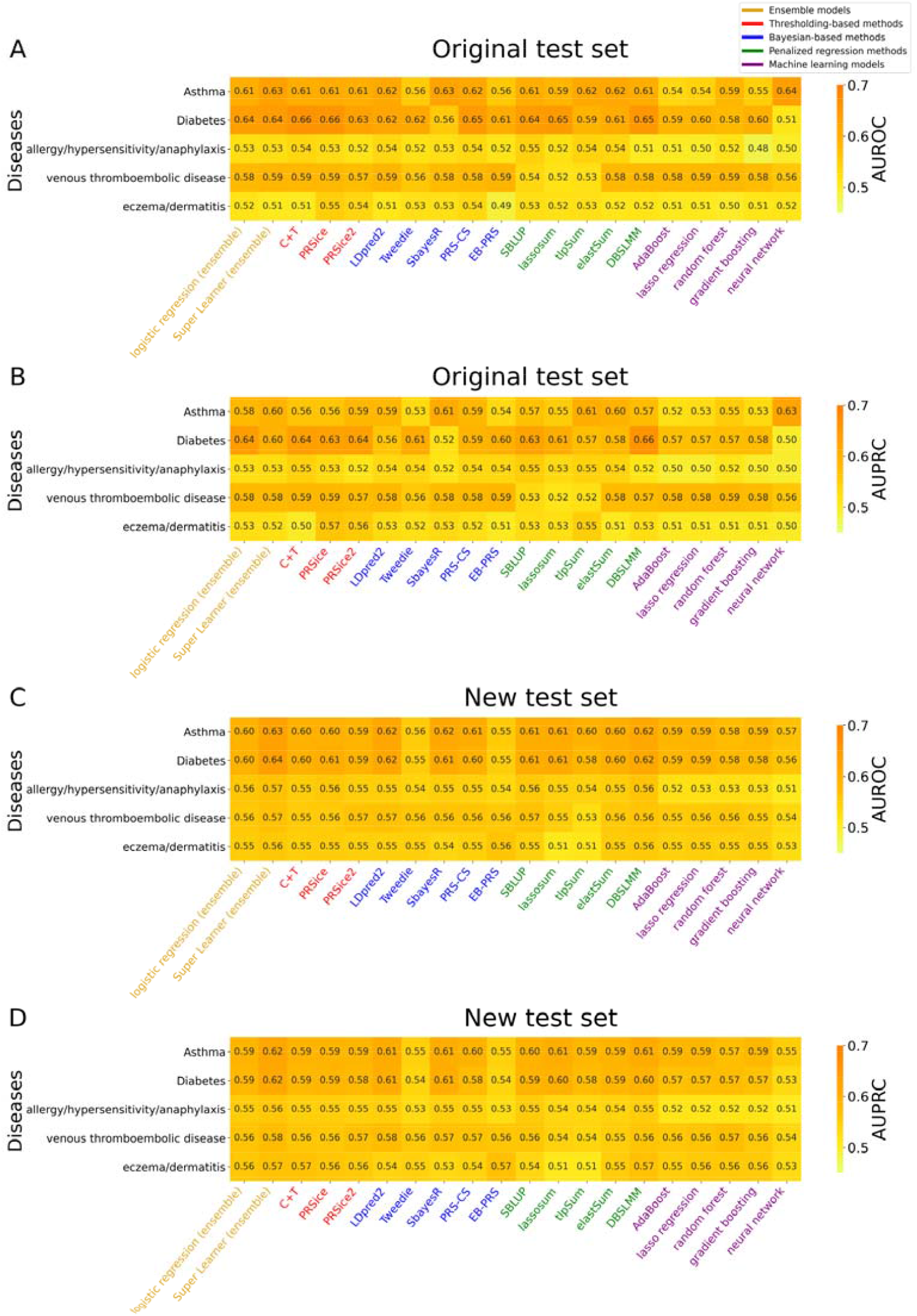
AUROC (**A** and **C**) and AUPRC (**B** and **D**) values of the method results on the five selected diseases from. The samples in the original test set are from incident cases, while the new test sets are derived from the validation set. The AUROC and AUPRC values for all the twenty-two diseases in are shown in Additional file 2: Figure S2.

**Figure 5.**
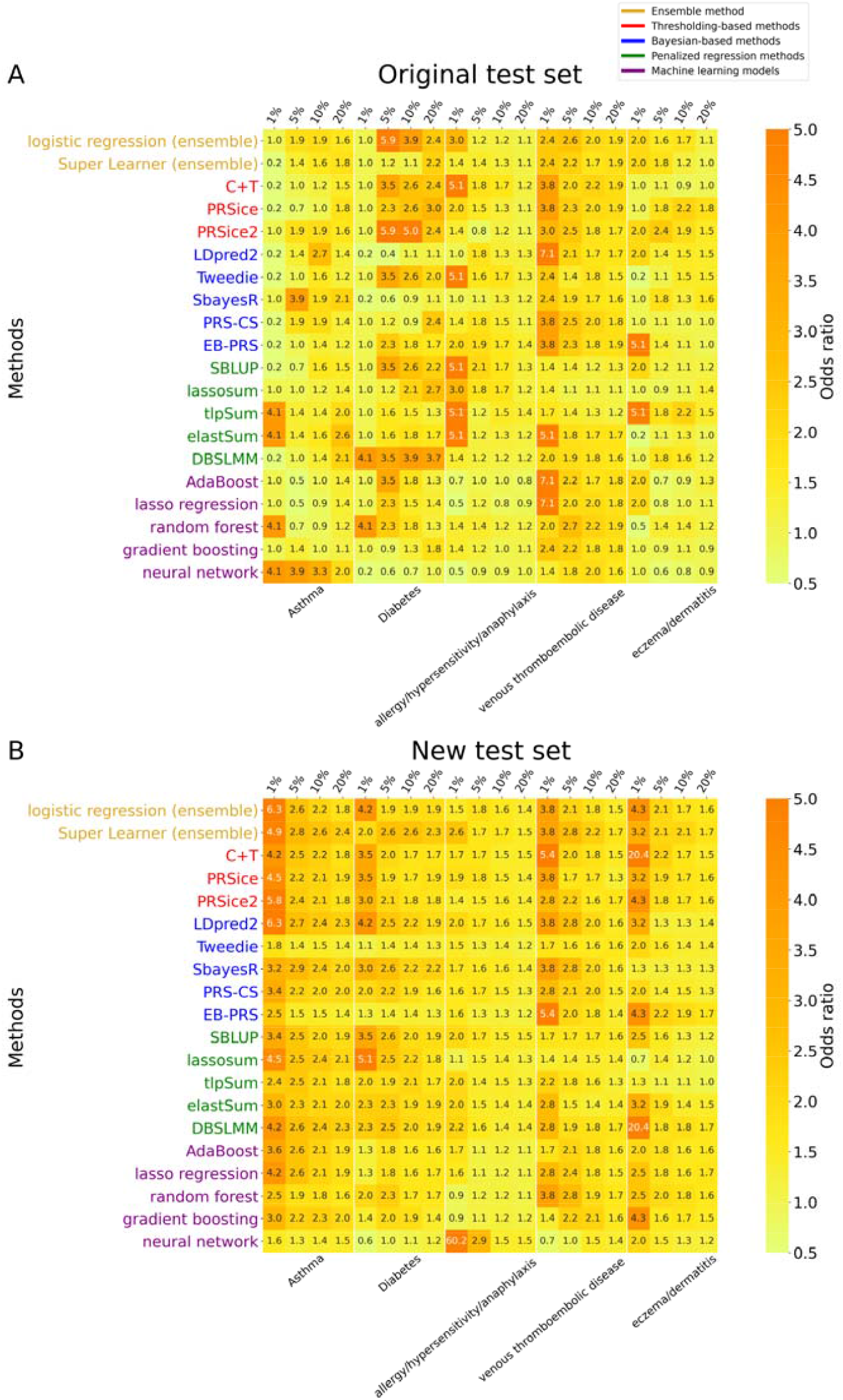
The ORs for the top 1, 5, 10 and 20 percentiles of the methods on the five selected diseases from. The samples in the original test set are from incident cases, while the new test sets are derived from the validation set. The OR values for all the twenty-two diseases in can be found in Additional file 4: Figure S4.

On the new test sets of prevalent cases, the overall performance of the methods is generally comparable to that when test sets of incident cases are used (Additional file 2: Figure S2C, S2D and Additional file 3: Figure S3B), and the ORs appear more realistic (Additional file 4: Figure S4B). Although the methods with good predictive performance tend to have greater power to identify high-risk individuals and achieve better ORs, the relatively poor predictive performance does not necessarily translate to poorer ORs. For instance, while the AUROCs of EB-PRS and C+T for venous thromboembolic disease are only 0.56 and 0.55 (Figure 4C), the top 1 percentile of ORs is 5.4 for both these methods (Figure 5B). We also investigated the circumstances that accounted for three extreme OR values in the top 1 percentile (Figure 5B): in neural network (OR = 60.2) for “allergy/hypersensitivity/anaphylaxis” and C+T (OR = 20.4) and DBSLMM (OR = 20.4) for “eczema/dermatitis”. For “allergy/hypersensitivity/anaphylaxis”, we found that as the neural network generated 39% individuals who had identical scores (51% of these were patients), the randomness of selecting individuals led to unstable ORs. For “eczema/dermatitis”, we found that the number of individuals in the top 1 percentile (i.e., 10 participants) was too small to achieve stable ORs for C+T and DBSLMM.

### Methods comparison using real data: predicting four common diseases with external summary statistics

The predictive performance for both statistical methods and machine learning models could be substantially influenced by the summary statistics (e.g. p-value, ORs, minor allele frequency et al.) of the training set. Many GWAS with very large sample sizes have enabled their summary statistics to be publicly available, which could be used as the training sets of statistical methods. We defined the patients of four diseases, including breast cancer (BC), coronary artery disease (CAD), inflammatory bowel disease (IBD), and type 2 diabetes (T2D), based on the descriptions in Khera et al.[31] using their self-reports and ICD-10 codes from hospitalization medical records (Additional file 7: Table S2 and Additional file 8: Table S3). The summary statistics of the four large cohort GWAS were downloaded from GWAS catalog [1, 32-34] for the corresponding diseases (Additional file 9: Table S4). We could not find any single method consistently better than the others, but some methods are stable to stay in the top 5 best in most cases, such as LDpred2 and PRS-CS. The methods, such as C+T, tlpSum and elastSum are significantly worse than other tools in all four diseases (Figure 6 and Additional file 5: Figure S5). When we considered the ORs in the top 1 percentile of PRSs of statistical and machine learning models, we found no tool was shown substantially better than the other ones for BC and IBD; For CAD and T2D, LDpred2, lassosum, DBSLMM and PRS-CS are generally more powerful than other tools (Figure 7).

**Figure 6.**
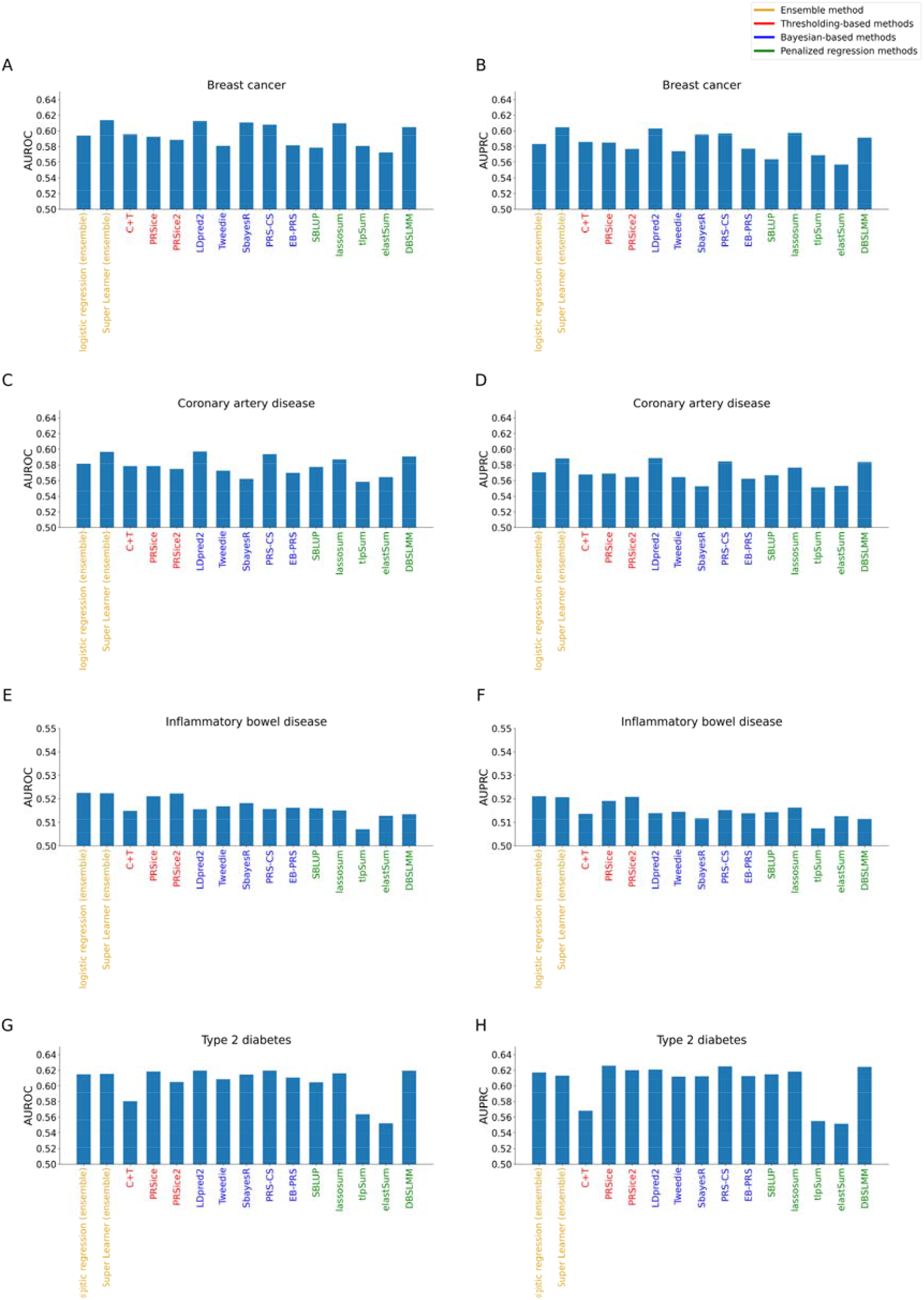
AUROC (**A, C, E, G**) and AUPRC (**B, D, F, H**) values of the statistical methods and ensemble models on the four diseases from.

**Figure 7.**
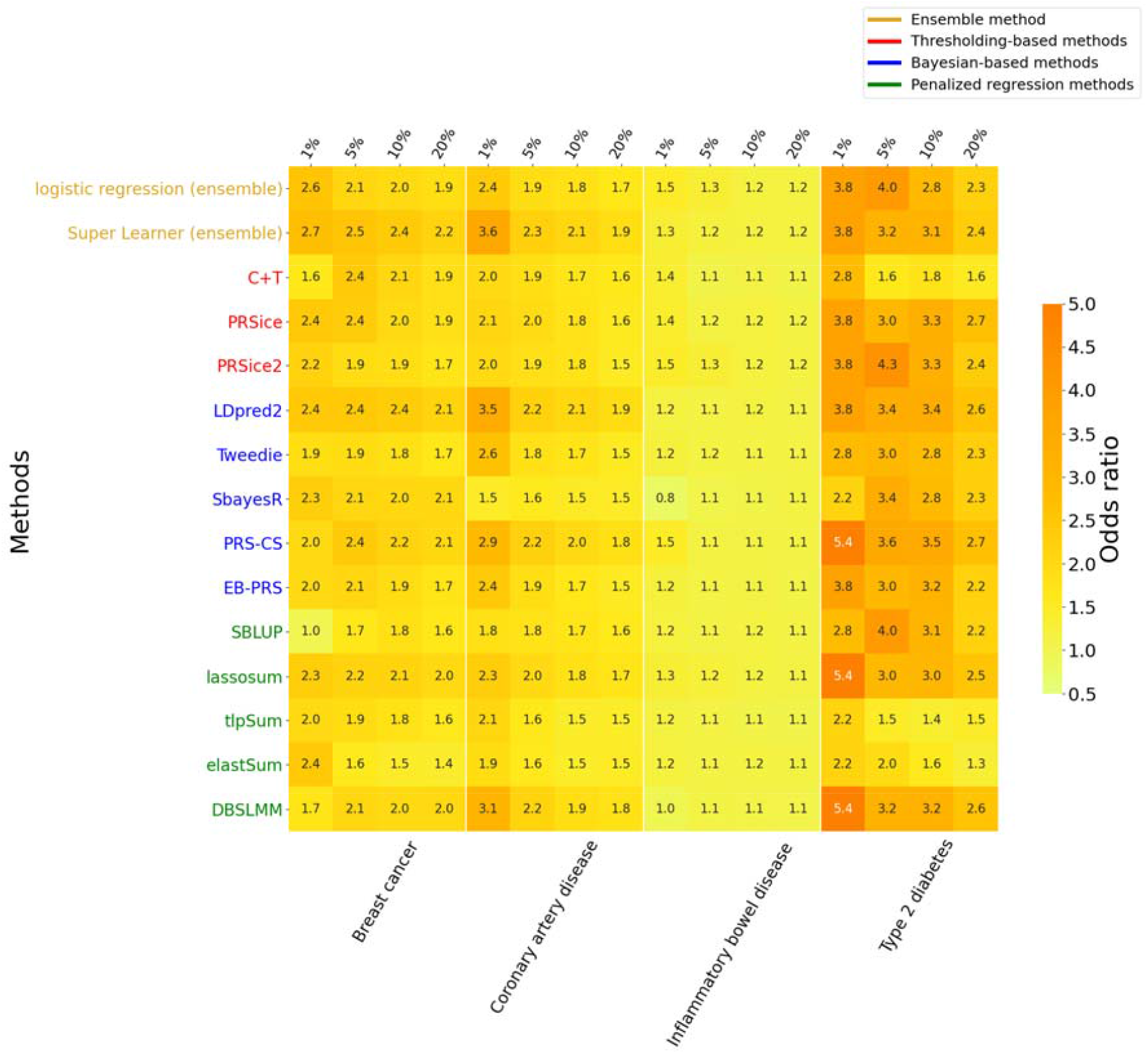
The ORs for the top 1, 5, 10, 20 percentiles of the statistical methods and ensemble models on the four diseases from.

### Ensemble models were stable and close to the best methods

We generated two ensemble models by integrating the results from thirteen statistical methods in logistic regression and Super Leaner (**Methods**). Super Learner is the better model compared to any individual method and the other ensemble model using logistic regression on the simulated data (Figure 2). It can also generate the most outstanding ORs even if the sample size is limited or disease heritability or SNP effect sizes are low (Figure 3). We observed a similar trend in the disease prediction of ensemble models for both *D*_22_ and *D*_4_, where Super Learner is always the best choice. Super Learner performs the best in more than eleven diseases in *D*_22_ and it is close to the best model in the remaining diseases (Additional file 2: Figure S2 and Additional file 3: Figure S3). Super Learner is one of the best tools in predicting BC, CAD and is close to the best model in IBD and T2D for *D*_4_ (Figure 6).

### Correlations between the methods

We calculated Spearman’s rank correlation coefficients (SRCCs) between the methods on *D*_22_ and *D*_4_ to investigate their correlations. We revealed machine learning models had higher SRCCs with each other than with other statistical or ensemble methods in *D*_22_ (Figure 8A). For statistical methods in *D*_*22*_, thresholding-based methods have higher SRCC (averaged SRCC = 0.74) than penalized regression methods (averaged SRCC = 0.63) and Bayesian-based methods (averaged SRCC = 0.54). A similar trend of the SRCCs between methods has been shown in *D*_4_, where the standard deviations become smaller (Figure 8B), suggesting the performance of statistical methods is more stable if the summary statistics are from large-cohort GWAS.

**Figure 8.**
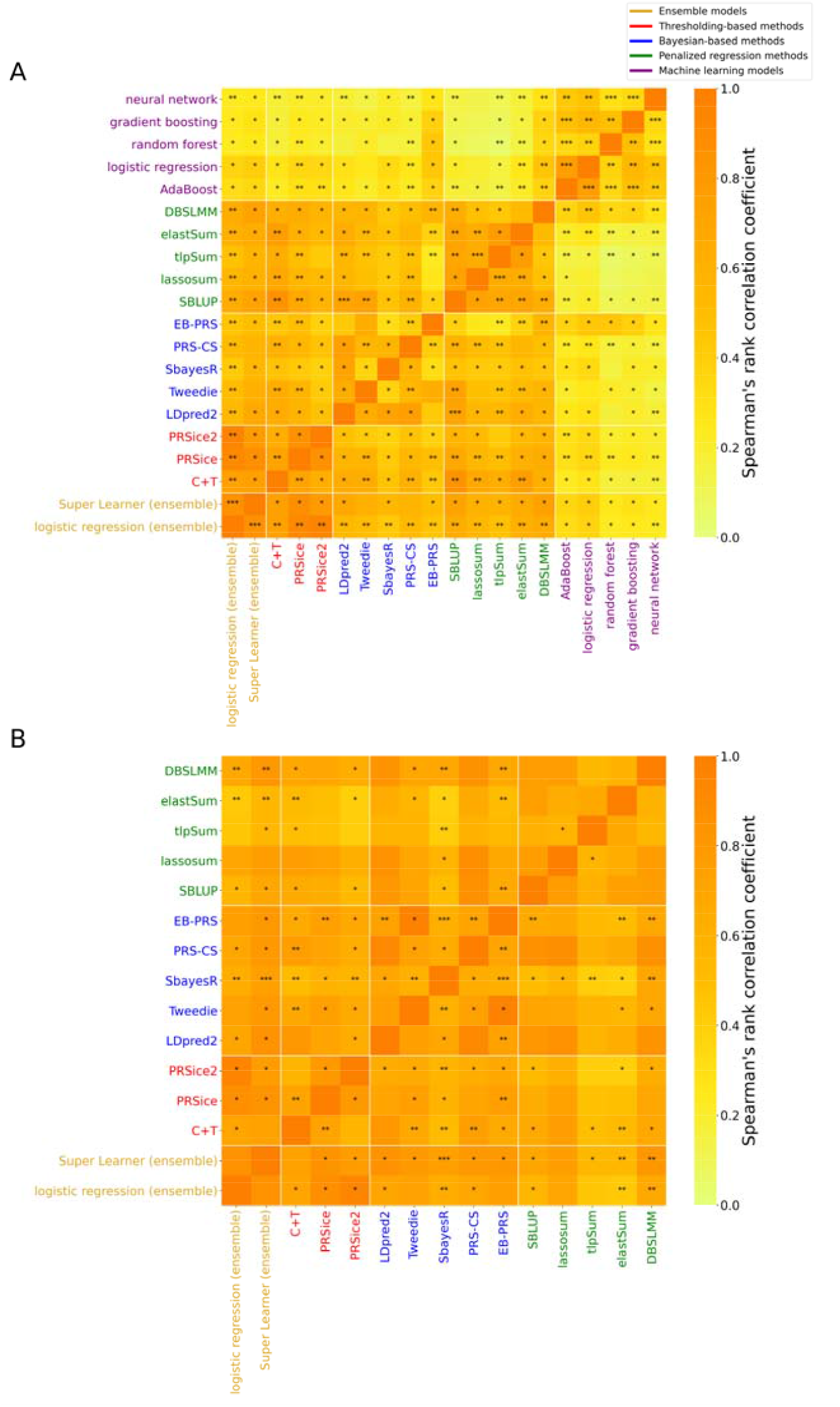
The average Spearman’s rank correlation coefficients between the methods based on the twenty-two diseases from (**A**) and four diseases from (**B**). The standard deviations are denoted as ‘***’ (larger than 0.3), ‘**’ (between 0.2 and 0.3), ‘*’ (between 0.1 and 0.2) and ‘ ’ (no larger than 0.1).

## Discussion

Advanced sequencing technologies have tremendously promoted research into the genetic factors for common diseases. As of July 8, 2021, the GWAS Catalog has collected 167,667 risk SNPs from 5,183 publications; many GWAS with extremely large sample sizes were also recently published in a blueprint of the common variants for particular diseases [35-38]. These studies motivated us to take advantage of well-documented genetic risk variants to improve the prediagnosis of clinical illnesses. It is particularly important to identify individuals in the general population who, owing to genetic factors, have increased risks for diseases, so as to facilitate early monitoring and treatment. The PRS has been the standard approach for linearly aggregating risk alleles and predicting disease risks. A variety of algorithms have been developed to implement PRS; however, a consensus on the best approach has not been reached.

In this study, we reviewed and analyzed thirteen statistical methods, five machine learning models and two ensemble models for PRS calculation and disease prediction. We observed several interesting trends that could be used to guide further algorithm development and model selection in clinical applications. First, GCTA [39] was used to generate the simulated datasets by an additive linear model. We identified the “simple” thresholding-based methods, such as C+T, PRSice and others, were often the best choices if the relationship between SNPs and the target disease was linear and all machine learning models were not good enough in such situations. This could be explained by their use of an insufficient sample size for learning a large number of parameters and the extra noise was introduced through the involvement of more irrelevant SNPs. The situation becomes more complex in real data and thresholding-based methods are no longer always better than other statistical methods. Second, the performance of PRS is significantly influenced by disease heritability. Diseases with high heritability are often more easily predicted across all methods. Third, both the numbers and effect sizes of risk SNPs are vital for distinguishing patients from controls. That is, if the number of association signals is sufficient—even when none of them is genome-wide significant (p-value < 5×10^−8^)—there would still be a possibility of identifying high-risk individuals with confidence (Figure 2C–D). Fourth, although the sample size of training set significantly influences the performance of machine learning models, it does not substantially affect the performance of most statistical methods. Fifth, the sophisticated ensemble models are promising in disease prediction and their performance is stable in different situations.

The PRSs of *D*_22_ from the UK Biobank data are found to be insufficient for application to clinical diagnosis (Addition file 2: Figure S2 and Additional file 3: Figure S3). This is possibly due to the strategy used for selecting patients in this study (**Methods**). To obtain sufficient sample sizes for training and validating the machine learning models, we focused on the most prevalent diseases in the UK Biobank. However, as disease status was assessed via participants’ self-reports and hospital ICD codes, our strategy for selecting patients may have conflated multiple subtypes of disorders. To clarify this issue, we selected four diseases with more precise definitions and used the external summary statistics from GWAS with large sample sizes. The results are not very optimistic and the performance of given tools are just slightly better than the random, such as IBD (Figure 6). We believe the power of PRS is more decisive by the inherent characteristics of diseases.

Population differences represent another factor that could potentially influence the PRS in clinical applications. All of the genotype data used in this study were from white British UK Biobank participants, and the conclusions were not affected by different characteristics of the populations (e.g., LD structures, disease locus heterogeneity). In clinical applications of PRS, cross-population trait predictions are usually underpowered and require careful monitoring. As the majority of GWAS were performed on populations of European descent, their findings could not be applied to other populations directly, due to the divergence in the minor allele frequency, LD structures and diseases locus heterogeneity. A recent study [40] showed that several common risk variants in a European-based GWAS were rarely found in East Asian and African populations, revealing key differences in the demographic histories of these populations. The effect sizes of risk alleles are determined by their local LD structures with the causal variants. As such, the diversity of LD structures may result in misleading PRS outcomes, especially when all risk SNPs must be aggregated across the entire human genome. For instance, when European-based effect sizes of the risk alleles were applied, prediction accuracies were reported to be 1.6-fold, 1.6-fold, 2.0-fold and 4.5-fold lower in Hispanic/Latino Americans, South Asians, East Asians and Africans, respectively [40]. Unfortunately, 49% of the studies and 78% of the samples published in the GWAS [41] are from populations of European ancestry, thus limiting the generalizability of PRS. Although several computational models have been developed to deal with this challenge [42, 43], more efforts should be dedicated to demonstrating their effectiveness.

In addition to genotype data used in the current study, other types of data can also provide power for predicting disease risk. For instance, Dogan et al. applied random forest to integrate genetic, epigenetic and phenotypic data from 1,545 individuals to predict the risk of CAD[44]. Xu et al.[45] constructed an online tool to predict irritable bowel syndrome from metagenomic datasets. Rather than using a single data type, a previous study used dynamic changes in personal integrative omics data to identify individuals’ medical risks for a diverse suite of diseases [46]. An important new direction in disease prediction is therefore to integrate multi-omics data with non-genetic factors (e.g. medical images, lifestyles and environmental factors) to jointly model common diseases.

## Conclusion

In this study, we benchmarked thirteen statistical methods, five machine learning and two ensemble models on simulated data, twenty-two common diseases with internal training sets and four diseases with external summary statistics from the UK Biobank. The effects of three key factors, including disease heritability, SNP effect size and sample size, on disease prediction are evaluated using simulated data. We also investigated the correlations between these methods and their standard deviations using the diseases defined by UK Biobank. We found the statistical methods were generally better than the machine learning models, suggesting the current sample size in training set did not support these complex machine learning models to capture both linear and nonlinear correlation between SNPs. Each method has its advantages and limitations and could be with substantially unstable performance for different diseases. We identified that disease heritability had a substantial effect on the predictive performance for all the methods. The number and effect sizes of risk SNP are equally important; the sample size in training set only strongly influences the performance of machine learning models. We also observed that the correlations of a pair of methods from the same category were relatively high. The external summary statistics from large cohort GWAS could reduce the standard deviation of the method correlations. We recommend to adopt a sophisticated ensemble model Super Learner, which is better than individual methods and generalized linear ensemble model for most cases.

## Method

### Preprocessing of genotype data from the UK Biobank

We performed quality control on 784,256 SNPs from 488,377 participants to remove non-informative and low-quality samples and SNPs. A total of 150,841 individuals were eliminated from the initial dataset if anyone satisfied at least one of the following criteria: (i) their self-reported ancestries were not “White British” (so as to control for population effects); (ii) they had more than 10% SNPs with missing genotypes; (iii) they were “outliers” or related samples that were excluded from principal component calculations; and (iv) they chose to withdraw their data. In addition, 287,679 SNPs were removed for failing to meet at least one of the following criteria: (i) their minor allele frequencies were lower than 1%; (ii) their p-values of Hardy–Weinberg equilibrium were less than 1 × 10^−6^; and

(iii) they showed genotype missing rates that exceeded 10%. The SNPs that passed the quality control were coded as 0, 1 and 2 according to the number of SNP minor alleles.

### Simulated data with varying disease heritability, SNP effect sizes and sample sizes

Our simulation was built on the processed UK Biobank genotype data after quality control; this consisted of 337,536 participants without disease labels and 39,420 SNPs from chromosome 1. We used GCTA[39] to generate the disease labels for the training, validation and test sets; this was done by varying disease heritability, SNP effect sizes and sample sizes (Figure 1, Table 2 and Additional file 13: Supplementary Note). We defined three random variables to represent “small”, “moderate” and “high” SNP effect sizes; these were drawn from normal distributions with means of 0 and variances of 0.01 (“small”), 0.1 (“moderate”) and 1 (“high”), respectively. The prevalence of disease was fixed to 0.2. The specific configurations of the simulated datasets are shown in Table 2. Each configuration was simulated 10 times so that the consistency and performance for each model could be examined.

### Patient definition and dataset construction for diseases from the UK Biobank data with internal training sets

Disease labels were identified based on self-reports and hospitalization medical records. Prevalent cases were defined as participants having self-reported diseases (including cancers) that were diagnosed before the baseline. The labels of self-reported diseases were organized using a “tree structure”. We considered the labels from the second level of the tree to guarantee that a sufficient number of patients were included. Data from hospitalization medical records included the ICD-10 codes of diseases and participants’ dates of diagnosis from hospital episodes. We generated a mapping table to match the ICD-10 codes with participants’ self-reported diseases based on the descriptions of their symptoms (Additional file 10: Table S5). The incident cases were defined as disease events occurring after the baseline (among participants who did not have prior disease events). Any male patients affected by breast cancer or gynecological disorders were removed. Any disease with fewer than 6,000 prevalent cases and 100 incident cases was eliminated. We generated a general control pool for participants who were never diagnosed with any of the investigated diseases. The controls were selected randomly and analyzed in relation to prevalent and incident cases. We used two-thirds of the prevalent cases and controls to generate the training sets, and the remaining one-third to generate the validation sets. The control samples were nonoverlapping between the training and validation sets. We collected incident cases and controls as test sets for investigating the predictive power of various models. We used the same number of controls as the cases to ensure the data was balanced for each disease.

### Patient definition and dataset construction for the four diseases from the UK Biobank data with external summary statistics

We also defined BC, CAD, IBD, and T2D using the similar criteria described in Khera et al.[31] (Additional file 7: Table S2) and collected their external GWAS summary statistics[1, 32-34] (Additional file 9: Table S4) to further evaluate the PRS models. The male patients with breast cancer were removed. According to summary statistics, the quality control on the SNPs was performed following the same criteria described in Privé et al.[11]. The remaining SNPs were kept if they belonged to both the UK Biobank and HapMap3[14]. The validation and test sets (Additional file 8: Table S3) were constructed based on the same strategy used to define the twenty-two common diseases, except that all the prevalent cases were included in the validation sets.

### Statistical methods for disease prediction

The GWAS summary statistics were either downloaded from publicly available GWAS studies for *D*_4_ or generated by logistic regression[47] on the training sets with age, gender and top 10 principal components as covariates for *D*_22_. They were used as inputs for the thirteen statistical methods (Table 1), including three thresholding-based methods, five Bayesian-based methods and five penalized regression methods. The software version and command lines for each method were included in Additional file 13: Supplementary Note. The reference panels for LD calculations were downloaded from the websites recommended by individual software. The LD was calculated by the UK Biobank samples if the software did not suggest any reference panels. For each tool, the hyperparameters were selected with respect to the optimal AUROC on the validation set from several candidates (Additional file 11: Table S6).

### Machine learning models for disease prediction

We generated four candidate SNP datasets according to the association p-value thresholds of 5 × 10^−3^, 5 × 10^−4^, 5 × 10^−5^ and 5 × 10^−6^ from the training set and removed one SNP from a SNP-pair if they had strong LD (*r*^2^ > 0.5) by using --indep-pairwise 50 5 0.5 in PLINK[47]. We used one linear model (lasso regression[48, 49]) and four nonlinear models (random forest[21, 22], AdaBoost[50, 51], gradient boosting[24, 25] and neural network[52, 53]) for disease prediction, and compared their performance with that of the statistical methods. For each machine learning model, we trained multiple times with different hyperparameter choices and used the validation set to select the best one based on the AUROC. Details of each model and hyperparameters can be found in Additional file 12: Table S7. In the application of neural network, we experimented using feedforward networks (of two to four layers) trained with gradient descent with a learning rate of 0.001 and adopted ReLu as an activation function. We shuffled the samples in each iteration and chose 200 samples as the size of the minibatch for the weight optimization by Adam[54]. The selected models were then tested on the independent test sets.

### Ensemble models for disease prediction

We implemented two ensemble models. One is a logistic regression which takes the results from thirteen statistical methods as variables. Their regression coefficients were trained using the validation sets, and the model was applied to the test sets for prediction. The other one, called Super Learner[55-57], is by creating multiple machine learning models and each model integrates the results of thirteen statistical methods. Super Learner then estimates the best weights of these models based on their predictive performance on the validation set, and integrates them as the final ensemble model, which could then be used to make predictions on the test sets.

### Model evaluation

We evaluated the predictive power of individual models using the value of the AUROC, AUPRC, PCC and the risk in the ascending percentile of prediction scores (odds ratios, ORs). The participants were sorted descendingly and binned into 100 separate groups according to their PRSs or predicted disease probabilities from the statistical methods, machine learning models and ensemble models. The predicted risk and the observed incidence of disease for each bin were then calculated. The predicted risk was calculated as the average of the disease probabilities across all individuals in each bin. For a given percentile X, ORs were calculated by comparing the numbers of incidence cases with prediction scores higher or lower than X. The correlations between methods were calculated using Spearman’s rank correlation coefficient.

## Supplementary information

**Additional file 1: Figure S1**. Pearson correlation coefficients (PCCs) between the method results and the simulated phenotypes by varying disease heritability (**A**), SNP effect sizes (**B**) and sample sizes (**C**).

**Additional file 2: Figure S2**. AUROC (**A** and **C**) and AUPRC (**B** and **D**) values of the method results on the twenty-two diseases from *D*_*22*_. The samples in the original test set are from incident cases, while the new test sets are derived from the validation set.

**Additional file 3: Figure S3**. Pearson correlation coefficients (PCCs) between the method results and the twenty-two diseases from *D*_22_. The samples in the original test set (A) are from incident cases, while the new test set (B) are derived from the validation set.

**Additional file 4: Figure S4**. The ORs for the top 1, 5, 10 and 20 percentiles of the methods on the twenty-two diseases from *D*_22_. The samples in the original test set are from incident cases, while the new test sets are derived from the validation set.

**Additional file 5: Figure S5**. Pearson correlation coefficients (PCCs) between the method results and the four diseases from *D*_*4*_.

**Additional file 6: Table S1**. The number of prevalent and incident cases for the twenty-two diseases from the UK Biobank data with internal training sets (*D*_22_). Prevalent cases are the patients diagnosed of diseases based on self-reports before the baseline examination and are used for training and validation. The incident cases are the individuals who changed status from non-disease to disease after the initial data collection and are used as the test set.

**Additional file 7: Table S2**. The definition of the four diseases from *D*_4_ based on the self-reports and ICD-10 codes.

**Additional file 8: Table S3**. The number of cases and controls in the validation sets and test sets for *D*_*4*_.

**Additional file 9: Table S4**. The information of GWAS summary statistics for the four diseases from *D*_*4*_.

**Additional file 10: Table S5**. Matching table between ICD-10 codes and self-reported diseases for twenty-two diseases (Excel).

**Additional file 11: Table S6**. Hyperparameters for statistical methods.

**Additional file 12: Table S7**. Hyperparameters for machine learning models.

**Additional file 13: Supplementary Note**.

## Acknowledgments

This research conducted using the UK Biobank Resource under Application 60434. We would like to thank James Zou for his useful comments in the early stage of this project.

## Authors’ contributions

LZ conceived the study. CW, LZ and JZ performed the simulation and UK Biobank data pre-processing. CW, LZ, JZ and XZ analyzed the results. LZ, CW and JZ wrote the manuscript. All authors read and approved the final manuscript.

## Availability of data and materials

UK Biobank data is from https://www.ukbiobank.ac.uk/. GWAS summary statistics of breast cancer, coronary artery disease, inflammatory bowel disease and type 2 diabetes are available in GWAS Catalog with PubMed ID of 29059683, 26343387, 26192919 and 28566273. The source code is available at https://github.com/Chonghao98/Methods-comparisons-on-disease-prediction. Links of individual software can be found in Additional file 13.

## Ethics approval and consent to participate

Not applicable.

## Author details

^1^Department of Computer Science, Hong Kong Baptist University, Hong Kong SRA, China

^2^Eye Institute and Department of Ophthalmology, NHC Key Laboratory of Myopia (Fudan University), Eye & ENT Hospital, Fudan University, Shanghai, China

^3^Department of Biomedical Engineering, Vanderbilt University, Vanderbilt Place Nashville, 37235, TN, USA

^4^Institute for Research and Continuing Education, Hong Kong Baptist University, Shenzhen, China

