## Additional information for "A comprehensive investigation of statistical and machine learning approaches for predicting complex human diseases on genomic variants"

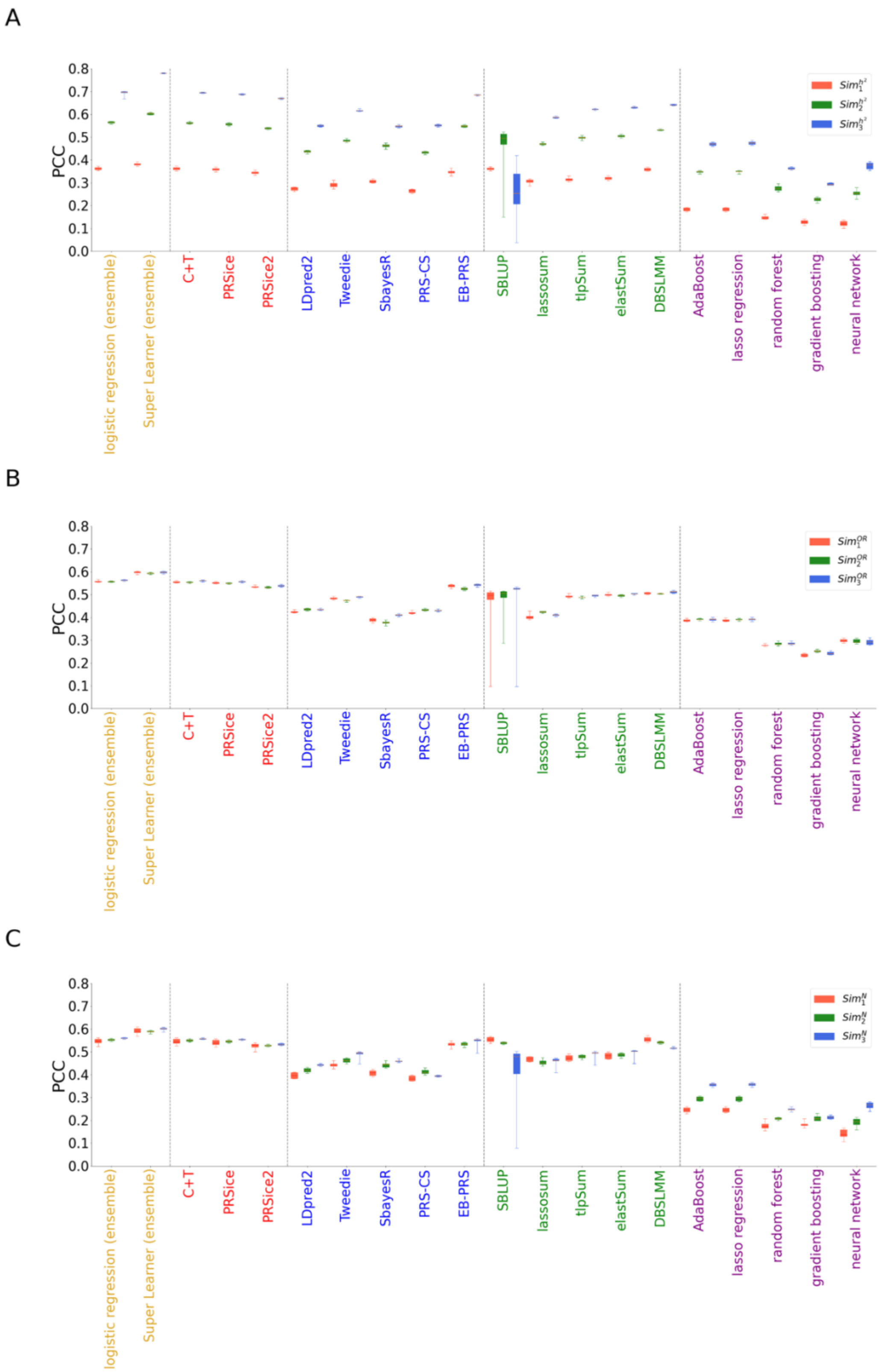


**Figure S1.** Pearson correlation coefficients (PCCs) between the method results and the simulated phenotypes by varying disease heritability (**A**), SNP effect sizes (**B**) and sample sizes (**C**).

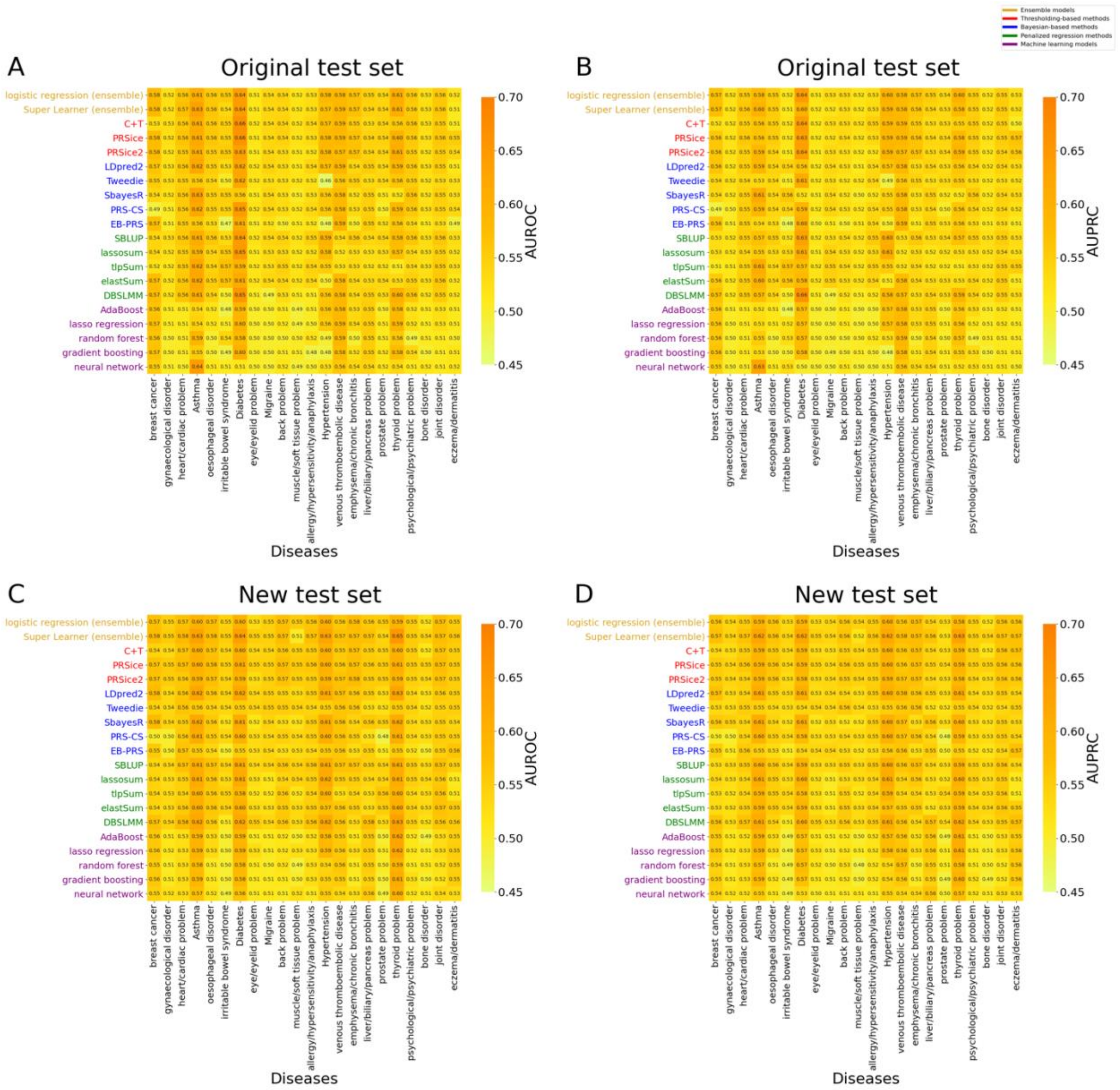


**Figure S2.** AUROC (**A** and **C**) and AUPRC (**B** and **D**) values of the method results on the twenty-two diseases from $D_{22}$. The samples in the original test set are from incident cases, while the new test sets are derived from the validation set.

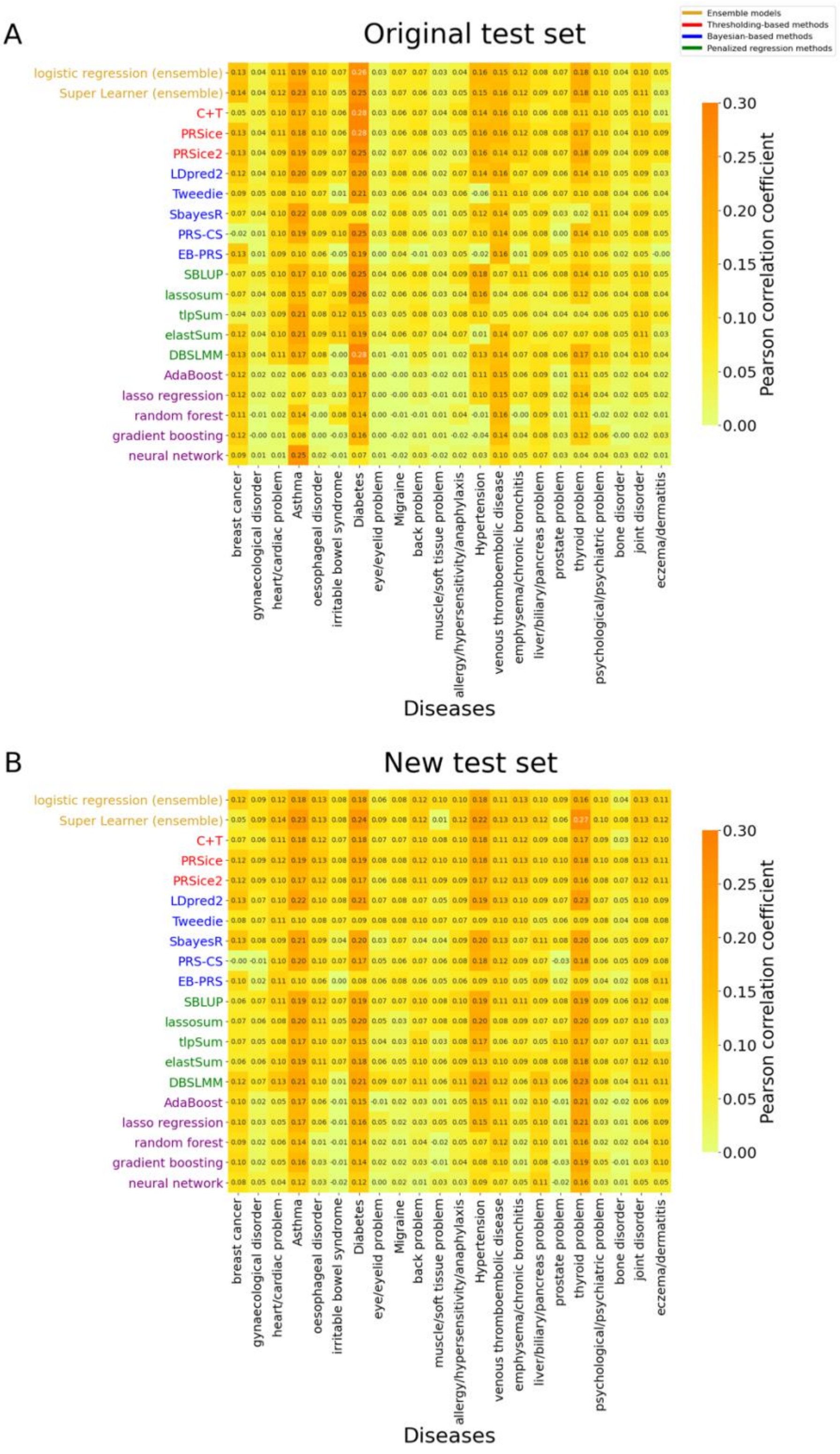


**Figure S3.** Pearson correlation coefficients (PCCs) between the method results and the twenty-two diseases from $D_{22}$. The samples in the original test set (**A**) are from incident cases, while the new test set (**B**) are derived from the validation set.

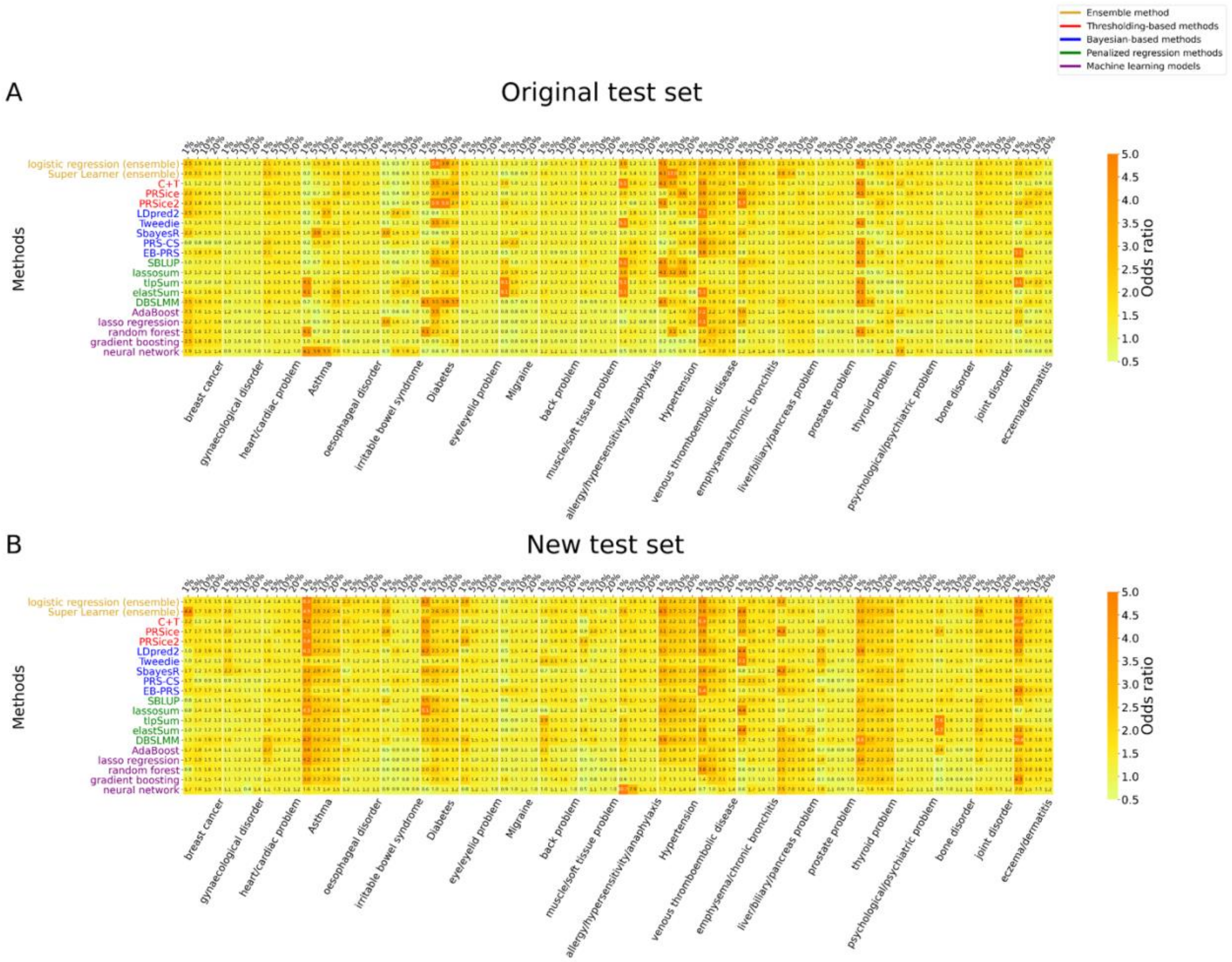


**Figure S4.** The ORs for the top 1, 5, 10 and 20 percentiles of the methods on the twenty-two diseases from $D_{22}$. The samples in the original test set are from incident cases, while the new test sets are derived from the validation set.

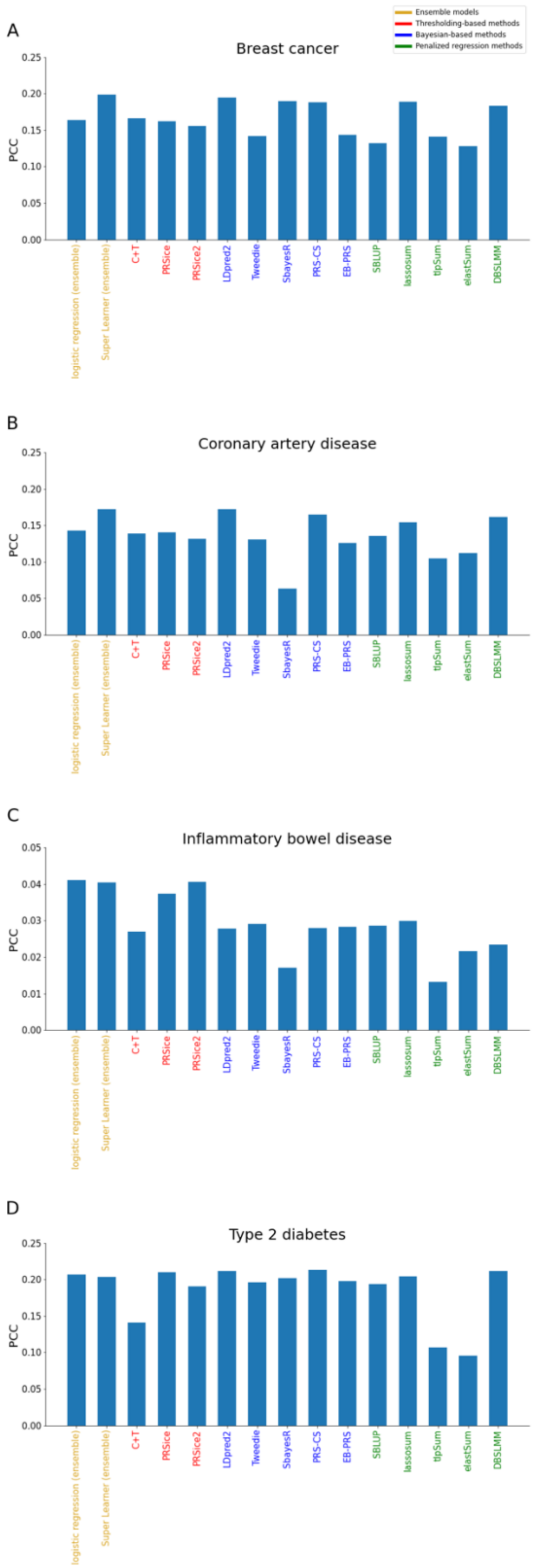


**Figure S5.** Pearson correlation coefficients (PCCs) between the method results and the four diseases from $D_4$.

| Disease Name | Prevalent cases | | | Incident cases | | |
|---|---|---|---|---|---|---|
| | Male | Female | Total | Male | Female | Total |
| Eczema/dermatitis | 4,401 | 5,162 | 9,563 | 137 | 195 | 332 |
| Gynaecological disorder | 0 | 16,446 | 16,446 | 0 | 7,273 | 7,273 |
| Allergy/hypersensitivity/anaphylaxis | 12,075 | 15,192 | 27,267 | 298 | 311 | 609 |
| Breast cancer | 0 | 7,641 | 7,641 | 0 | 3,176 | 3,176 |
| Migraine | 2,464 | 8,236 | 10,700 | 126 | 328 | 454 |
| Muscle/soft tissue problem | 3,701 | 4,290 | 7,991 | 2,376 | 1,960 | 4,336 |
| Bone disorder | 1,295 | 6,557 | 7,852 | 1,406 | 2,093 | 3,499 |
| Irritable bowel syndrome | 2,243 | 6,320 | 8,563 | 66 | 173 | 239 |
| Psychological/psychiatric problem | 9,494 | 17,195 | 26,689 | 551 | 419 | 970 |
| Thyroid problem | 3,147 | 16,889 | 20,036 | 26 | 114 | 140 |
| Asthma | 16,978 | 22,576 | 39,554 | 57 | 87 | 144 |
| Oesophageal disorder | 11,012 | 12,641 | 23,653 | 3,187 | 3,329 | 6,516 |
| Liver/biliary/pancreas problem | 3,291 | 6,475 | 9,766 | 2,252 | 3,567 | 5,819 |
| Back problem | 10,354 | 11,149 | 21,503 | 1,717 | 2,250 | 3,967 |
| Joint disorder | 19,183 | 26,109 | 45,292 | 4,560 | 5,870 | 10,430 |
| Eye/eyelid problem | 7,428 | 8,011 | 15,439 | 5,815 | 7,333 | 13,148 |
| Venous thromboembolic disease | 3,755 | 5,138 | 8,893 | 671 | 549 | 1,220 |
| Heart/cardiac problem | 16,186 | 8,850 | 25,036 | 2,847 | 1,640 | 4,487 |
| Diabetes | 10,277 | 5,961 | 16,238 | 98 | 41 | 139 |
| Prostate problem | 6,427 | 0 | 6,427 | 2,338 | 0 | 2,338 |
| Hypertension | 48,427 | 43,631 | 92,058 | 44 | 81 | 125 |
| Emphysema/chronic bronchitis | 3,747 | 3,794 | 7,541 | 577 | 445 | 1,022 |

**Table S1**. The number of prevalent and incident cases for the twenty-two diseases from the UK Biobank data with internal training sets ($D_{22}$). Prevalent cases are the patients diagnosed of diseases based on self-reports before the baseline examination and are used for training and validation. The incident cases are the individuals who changed status from non-disease to disease after the initial data collection and are used as the test set.

| Disease | Self-reports | ICD-10 codes |
|---|---|---|
| Breast cancer | In UKBB | C50.X |
| Coronary artery disease | In UKBB | I21.X, I22.X, I23.X, I24.1, I25.2 |
| Inflammatory bowel disease | In UKBB | K51.X |
| Type 2 diabetes | In UKBB | E11.X |

**Table S2.** The definition of the four diseases from $D_4$ based on the self-reports and ICD-10 codes.

| Disease | Validation set | | Test set | |
|---|---|---|---|---|
| | cases | controls | cases | controls |
| Breast cancer | 6615 | 6615 | 4649 | 4649 |
| Coronary artery disease | 9401 | 9401 | 11342 | 11342 |
| Inflammatory bowel disease | 8220 | 8220 | 18968 | 18968 |
| Type 2 diabetes | 2609 | 2609 | 965 | 965 |

**Table S3.** The number of cases and controls in the validation sets and test sets for $D_4$.

| Disease | Publication | Cases | Controls |
|---|---|---|---|
| Breast cancer | Michailidou et al.[1] | 122977 | 105974 |
| Coronary artery disease | Nikpay et al.[2] | 60801 | 123504 |
| Inflammatory bowel disease | Liu et al.[3] | 12882 | 21770 |
| Type 2 diabetes | Scott et al. [4] | 26676 | 132532 |

**Table S4.** The information of GWAS summary statistics for the four diseases from $D_4$.

| | | | | | | | | | | | | | | | | | |
|---|---|---|---|---|---|---|---|---|---|---|---|---|---|---|---|---|---|
| Eczema/dermatitis | L20-L30 | | | | | | | | | | | | | | | | |
| Gynaecological disorder | N97 | N83 | E28 | D25 | D26 | N84 | N81 | N80 | N87 | N86 | N92 | N73 | N74 | N94 | N95 | A56 | A54 |
| Allergy/hypersensitivity/anaphylaxis | T78 | T88 | J30 | L50 | L23 | Z88 | D69 | | | | | | | | | | |
| Breast cancer | C50 | | | | | | | | | | | | | | | | |
| Migraine | G43 | | | | | | | | | | | | | | | | |
| Muscle/soft tissue problem | M60 | G72 | M79 | M72 | | | | | | | | | | | | | |
| Bone disorder | M86 | M80 | M81 | M88 | M25 | | | | | | | | | | | | |
| Irritable bowel syndrome | K58 | | | | | | | | | | | | | | | | |
| Psychological/psychiatric problem | F01-F99 | | | | | | | | | | | | | | | | |
| Eye/eyelid problem | H00-H59 | | | | | | | | | | | | | | | | |
| Thyroid problem | E05 | E02 | E03 | E07 | | | | | | | | | | | | | |
| Asthma | J45 | J46 | | | | | | | | | | | | | | | |
| Oesophageal disorder | K20 | K21 | K22 | K23 | | | | | | | | | | | | | |
| Liver/biliary/pancreas problem | K70-K77 | K80-K87 | | | | | | | | | | | | | | | |
| Back problem | M45-M49 | M54 | | | | | | | | | | | | | | | |
| Joint disorder | M05 | M06 | M15 | M16 | M17 | M18 | M19 | M10 | M25 | M13 | | | | | | | |
| Venous thromboembolic disease | I26 | I81 | I82 | | | | | | | | | | | | | | |
| Heart/cardiac problem | I20 | I21 | I50 | J81 | I49 | R01 | I38 | I42 | I43 | I31 | I51 | I01 | | | | | |
| Diabetes | O24 | E10 | E11 | E12 | E13 | E14 | | | | | | | | | | | |
| Prostate problem | N40 | N41 | N42 | | | | | | | | | | | | | | |
| Hypertension | I10 | O13 | | | | | | | | | | | | | | | |
| Emphysema/chronic bronchitis | J43 | J44 | J98 | | | | | | | | | | | | | | |

**Table S5.** Matching table between  ICD 10 codes and self-reported diseases.

| Methods | Hyperparameters | Par1 | Par2 | Par3 | Par4 | Par5 | Par6 | Par7 |
|---|---|---|---|---|---|---|---|---|
| C+T | p | 1 | 0.5 | 0.05 | 5e-04 | 5e-06 | 5e-08 | - |
| | r2 | 0.2 | 0.4 | 0.6 | 0.8 | - | - | - |
| PRSice | --clump-p | 1 | 0.5 | 0.05 | 5e-04 | 5e-06 | 5e-08 | - |
| | --clump-r2 | 0.2 | 0.4 | 0.6 | 0.8 | - | - | - |
| PRSice2 | NA | - | - | - | - | - | - | - |
| LDpred2 | p | 1 | 0.3 | 0.1 | 0.03 | 0.01 | 0.003 | 0.001 |
| | $h^2$ | $0.7*h^2$_est | $1*h^2$_est | $1.4*h^2$_est | - | - | - | - |
| Tweedie | NA | - | - | - | - | - | - | - |
| SbayesR | chain-length | 5000 | 10000 | 50000 | - | - | - | - |
| | burn-in | 2000 | 10000 | 20000 | - | - | - | - |
| PRS-CS | Phi | 1 | 0.01 | 1e-04 | 1e-06 | - | - | - |
| EB-PRS | NA | - | - | - | - | - | - | - |
| SBLUP | cojo-wind | 1000 | - | - | - | - | - | - |
| lassosum | NA | - | - | - | - | - | - | - |
| tlpSum | lambda | 0.1 | 0.5 | 1 | - | - | - | - |
| | tau | 5e-03 | 1e-03 | 5e-04 | 1e-04 | 5e-05 | 1e-05 | 1e-06 |
| elastSum | lambda | 0.1 | 0.5 | 1 | - | - | - | - |
| | alpha | 0 | 0.5 | 1 | 1.5 | - | - | - |
| DBSLMM | pv | 1e-05 | 1e-06 | 1e-07 | 1e-08 | - | - | - |
| | r2 | 0.05 | 0.1 | 0.15 | 0.2 | 0.25 | - | - |

**Table S6.** Hyperparameters chosen for statistical methods. p: p value threshold; r2: $r^2$ threshold; clump-p: the p-value threshold used for clumping; clump-r2: the $r^2$ threshold for clumping; h2: disease heritability, h2_est: the estimated disease heritability; chain-length: the total number of iterations in MCMC; burn-in: the number of iterations to be discarded; phi: global shrinkage parameter; cojo-wind: specify a distance *d* (in Kb unit). LD between SNPs more than *d* Kb away from each other are ignored. Lambda, tau and alpha: tuning parameters in the formulas of tlpSum or elastSum; pv: p value threshold. Par1 to Par7 are the seven candidate values evaluated in the validation set.

| Model | Hyperparameter | Par1 | Par2 | Par3 | Par4 |
|---|---|---|---|---|---|
| lasso regression | regularization | 10 | 100 | 1000 | 10000 |
| random forest | min_impurity_decrease | 0.1 | 0.01 | 0.001 | 0.0001 |
| neural network | structure | (20,20) | (30,30) | (10,10,10,10) | (30,20,10) |
| adaboost | base_classifier | DT | LR | ET | GNB |
| gradient boosting | min_impurity_decrease | 0.1 | 0.01 | 0.001 | 0.0001 |

**Table S7.** Hyperparameters for training machine learning models. Regularization: regularization strength in Lasso Regression; min_impurity_decrease: a node would be split if the decrease of the impurity is no smaller than this value; structure: the number of hidden nodes for each layer (split by comma); base_classifier: base classifiers embedded in adaboost, DT: Decision Tree; LR: Logistic Regression; ET: Extra Tree; GNB: Gaussian Naïve Bayesian. Par1 to Par4 are the four candidate values evaluated in the validation set.

### Supplementary Note

#### Steps to generate simulated data

(a) Prepare genotype data (.bed, .bim and .fam PLINK files, with individuals' phenotypes are missing) and covariance file (age, gender, and top 10 principal components).

(b) Split the individuals in the .fam file into three parts by random to construct training, validation and test sets based on the given percentage. And split raw genotype accordingly by PLINK.

- plink --bfile prefix_of_bed_file (raw_bed_file) --keep txt_file (contains_individuals_of_interest) --make-bed --out prefix_of_output

(c) Select a list of SNPs to be risk variants, and assign their effect sizes.

(d) Determine disease heritability, disease prevalence, the numbers of cases and controls and the number of simulation replicates.

(e) Apply GCTA to generate the phenotypes for the individuals in the training set.

- gcta64 --bfile prefix_of_bed_file (training_set) --simu-cc number_of_cases number_of_controls --simu-causal-loci risk_SNP_list --simu-hsq heritability --simu-k disease_prevalence --simu-rep number_of_replicates --out prefix_of_output

(f) Generate the association statistics by PLINK on the training set.

- plink --bfile prefix_of_bed_file (training_set) --geno 0.1 --hwe 0.000001 --maf 0.01 --mind 0.1 --logistic (--covar covariance_file) --freq --ci 0.95 --out prefix_of_output

(g) Extract the effect sizes of all the SNPs from the summary statistics.

(h) Apply GCTA to generate the phenotypes for the validation set and test set.

- gcta64 --bfile prefix_of_bed_file (validation_set_or_test_set) --simu-cc number_of_cases number_of_controls --simu-causal-loci all_SNP_list --simu-hsq heritability --simu-k disease_prevalence --simu-rep number_of_replicates --out prefix_of_output

### Command lines, links and versions for each method

**1. C+T (version 1.0.10)**
Link: https://github.com/bvilhjal/ldpred
Command line:
ldpred p+t --cf coordinated_file --ldr LD_radius --p tuning_parameter --r2 tuning_parameter --out output

**2. PRSice (version 2.3.1.e)**
Link:
https://github.com/choishingwan/PRSice/tree/dc04caa38c05e9f15484edaeb1cfce341da7cf1d
Command line:
Rscript PRSice.R --prsice PRSice --base summary_statistics_file --target target_data --thread number_of_threads --clump-p tuning_parameter --clump-r2 tuning_parameter --out output

**3. PRSice2 (version 2.3.3)**
Link:
https://github.com/choishingwan/PRSice/tree/b075dda6c93b11ef1ba5ff81fb2fbaaff70ed9e3
Command line:
Rscript PRSice.R --prsice PRSice_linux --base summary_statistics_file --stat OR --or --target target_data --pheno phenotype_file --cov covariance_file --out output

**4. LDpred2 (bigsnpr version 1.6.1)**
Link: https://privefl.github.io/bigsnpr/articles/LDpred2.html
Command line:
Rscript LDpred2.R --bfile reference_panel --geneticPos 1000G_file --sst summary_statistics_file --ncores number_of_threads --out output
LDpred2.R is available in https://github.com/Chonghao98/Methods-comparisons-on-disease-prediction

**5. Tweedie (version 1)**
Link: https://sites.google.com/site/honcheongso/software/empirical-bayes-risk-prediction
The script is available in https://github.com/Chonghao98/Methods-comparisons-on-disease-prediction

**6. SbayesR (version 2.0)**
Link: https://cnsgenomics.com/software/gctb/#Download
Command line:
gctb --sbayes R --ldm LD_matrix --pi 0.95,0.02,0.02,0.01 --gamma 0,0.01,0.1,1 --gwas-summary summary_statistics_file --chain-length tuning_parameter --burn-in tuning_parameter --out output

**7. PRS-CS (the version is the one updated on April 24, 2020)**
Link: https://github.com/getian107/PRScs
Command line:
python3 PRScs.py --ref_dir=reference_panel --bim_prefix=bim_file_prefix --sst_file=summary_statistics_file --chrom=chromosome(s) --n_gwas=sample_size_of_gwas --out_dir=output --phi=tuning_parameter

**8. EB-PRS (version 2.1.0)**
Link: https://cran.r-project.org/web/packages/EBPRS/index.html
Command line:
Rscript EBPRS.R --sst summary_statistics_file --cc .frq.cc_file --out output
EBPRS.R is in https://github.com/Chonghao98/Methods-comparisons-on-disease-prediction

**9. SBLUP (version 1.93.2 beta)**
Link: https://cnsgenomics.com/software/gcta/#Download
Command line:
gcta64 --bfile prefix_of_bed_file --cojo-file summary_statistics_file --cojo-sblup lambda --cojo-wind LD_distance --thread-num number_of_threads --out output

**10. lassosum (version 0.4.5)**
Link: https://github.com/tshmak/lassosum
Command line:
Rscript lassosum.R --bfile reference_panel --sst summary_statistics_file --ncores number_of_threads --covariate covariance_file --out output
lassosum.R is available in https://github.com/Chonghao98/Methods-comparisons-on-disease-prediction

**11. tlpSum (version 0.0.0.9000)**
Link: https://github.com/jpattee/penRegSum
Command line
Rscript tlpSum.R --ref reference_panel --sst summary_statistics_file --lambda tuning_parameter --tau tuning_parameter --out output
tlpSum.R is available in https://github.com/Chonghao98/Methods-comparisons-on-disease-prediction

**12. elastSum (version 0.0.0.9000)**
Link: https://github.com/jpattee/penRegSum
Command line:
Rscript elastSum.R --ref reference_panel --sst summary_statistics_file --lambda tuning_parameter --alpha tuning_parameter --out output
elastSum.R is available in https://github.com/Chonghao98/Methods-comparisons-on-disease-prediction

**13. DBSLMM (version 0.1)**
Link:
https://github.com/biostat0903/DBSLMM/tree/cf4d494dfe75a6bc911265cb150059898502a80f
Command line:
Rscript DBSLMM.R --summary summary_statistics_file --type t --outPath output_path --plink plink --dbslmm dbslmm --ref reference_panel --pv p_value_threshold --r2 r2_threshold --mafMax 1 --n sample_size_of_summary_data --nsnp the_number_of_SNPs --block block_information --h2 heritability

**14. adaboost (scikit-learn version 0.22.1)**
Link:
https://scikit-learn.org/stable/modules/generated/sklearn.ensemble.AdaBoostClassifier.html
The script is in https://github.com/Chonghao98/Methods-comparisons-on-disease-prediction

**15. lasso regression (scikit-learn version 0.22.1)**
Link:
https://scikit-learn.org/stable/modules/generated/sklearn.linear_model.LogisticRegression.html
The script is available in https://github.com/Chonghao98/Methods-comparisons-on-disease-prediction

**16. random forests (scikit-learn version 0.22.1)**

Link:
https://scikit-learn.org/stable/modules/generated/sklearn.ensemble.RandomForestClassifier.html
The script is available in https://github.com/Chonghao98/Methods-comparisons-on-disease-prediction

**17. gradient boosting (scikit-learn version 0.22.1)**
Link:
https://scikit-learn.org/stable/modules/generated/sklearn.ensemble.GradientBoostingClassifier.html
The script is available in https://github.com/Chonghao98/Methods-comparisons-on-disease-prediction

**18. neural network (scikit-learn version 0.22.1)**
Link:
https://scikit-learn.org/stable/modules/generated/sklearn.neural_network.MLPClassifier.html
The script is available in https://github.com/Chonghao98/Methods-comparisons-on-disease-prediction

**19. logistic regression ensemble model**
The script is available in https://github.com/Chonghao98/Methods-comparisons-on-disease-prediction

**20. Super Learner ensemble model (version 2.0-28)**
Link:
https://cran.r-project.org/web/packages/SuperLearner/index.html
The script is available in https://github.com/Chonghao98/Methods-comparisons-on-disease-prediction
